# Does plant root architecture respond to potassium nutrition under water stress? A case from rice seedling root responses

**DOI:** 10.1101/2020.08.05.237685

**Authors:** Dipika S. Patel, Bardhan Kirti, P Patel Dhiraji, Parekh Vipulkumar, Jena Suchismita, V. Narwade Ajay, Harshadkumar N Chhatrola

## Abstract

The root is the sensing organ for potassium (K) and water availability. We evaluated whether K availability influences root architecture and contributes to drought tolerance under moisture stress. Rice seedling growth was severely affected by low K availability under water stress, and the substantial reductions in root projected area, maximum width, and width to depth ratio were observed. High K availability helps maintain root top and bottom angles and reduces root steepness under mild water stress, but over K nutrition does not ensure higher seedling growth. Under severe water stress, the steepness was more regulated by water than K availability.

## INTRODUCTION

In rice, root thickness, density, depth, and distribution have been considered as key traits contributing to drought tolerance, depending on the target environment^1^. Early-stage changes in root angle, which develop deep root systems, contribute to drought tolerance in rice^2^. Growing evidence suggests a cross-talk between mechanisms for water and nutrient sensing in roots; and root signals modulate the shoot growth in response to water and nutrient availability^3^. Potassium (K) fertilization is a common strategy for alleviating the negative effects of water stress in many crops, including rice^4,5^. High potassium level increases root diameter and dry matter^6^. The effect of K on the root is attributed to hormonal balance^7^. Briefly, the literature suggests that in addition to playing a key role in maintaining water balance in cells^8^, regulating stomatal conductance^9^ and neutralizing reactive oxygen species^10^, K might also influence root growth and development in a three-dimensional space, termed as root architecture, during moisture stress. There is a gap in the knowledge on the influence of K on root architecture under moisture stress, as most studies either focused on root growth^11^ or were conducted under no water stress^6^. Similarly, the influence of water availability on rice root system has been studied^12^; however, very little is known about the combined effects of water and nutrient availability. The main objective of the present study was to evaluate the effect of K on root architecture and growth of rice seedlings under moisture stress.

## MATERIALS AND METHODS

NAUR-1, a variety of rice (*Oryza sativa* L.), has wide adaptability to upland and lowland cultivation in the south Gujarat region. The experiment was conducted under a naturally ventilated polyhouse at the institute (20.9248° N, 72.9079° E) during 2017–18 in a completely randomized factorial design with three replications. The seeds were pre-soaked in deionized water for 24 h, and five seeds were sown per semi-transparent polythene bag (32 cm x 20.5 cm) containing 1.2 Kg of a uniform mixture of sand and perlite (2:1, 800 g +400 g). After emergence, only two seedlings were retained per bag.

The experiment consisted of three water stress levels, including NWS (No water stress), MWS (mild water stress), and SWS (severe water stress), corresponding to 100, 60, and 40% of field capacity (FC), respectively. The gravimetric method was used to estimate FC. Four water-saturated bags, with covered top to prevent evaporation, were kept overnight to drain the excess water. To calculate the amount of water retained at 100% of the FC, the following formula was used: [(the weight of saturated media – the weight of oven-dried media) / the weight of oven-dried media] × 100; media were kept in a hot air oven at 105 °C ±1 °C overnight to dry to a constant weight. Daily evapotranspiration was estimated by daily weighing of four bags, and their weight differences with 100% of FC were recorded. Initially, all bags were daily replenished by the unmodified Yoshida solution^13^ to maintain 100% FC until the seedling emergence, and thereafter, the modified Yoshida solution was used to simulate the water stress (Figure 1 A, B, and Figure 2).

**Figure 1.**
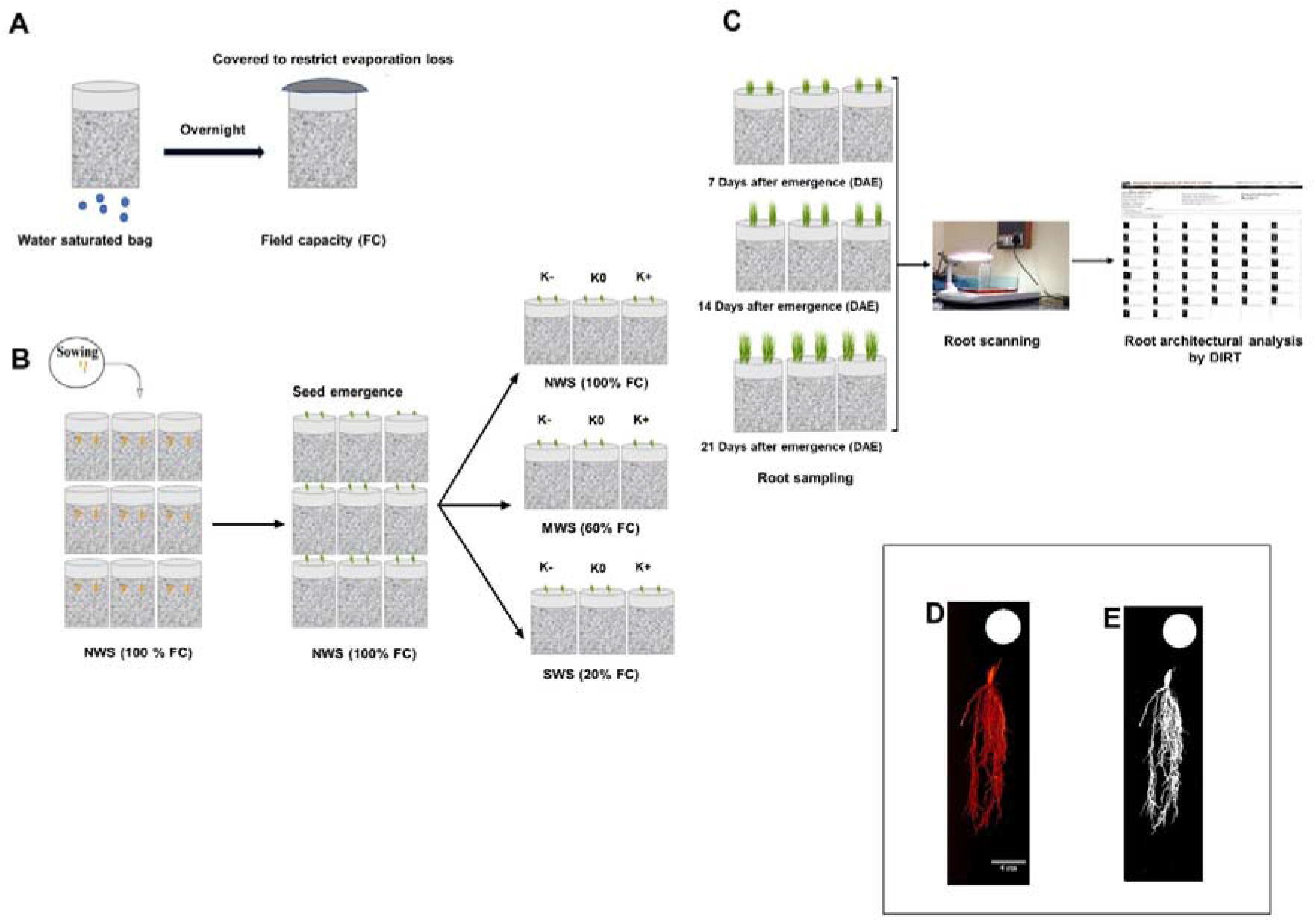
Scheme of experimental setup: estimation of field capacity (A), treatment imposition (B), analysis of root architecture (C). Inset: scanned root image (D) and masked root image (E).

**Figure 2.**
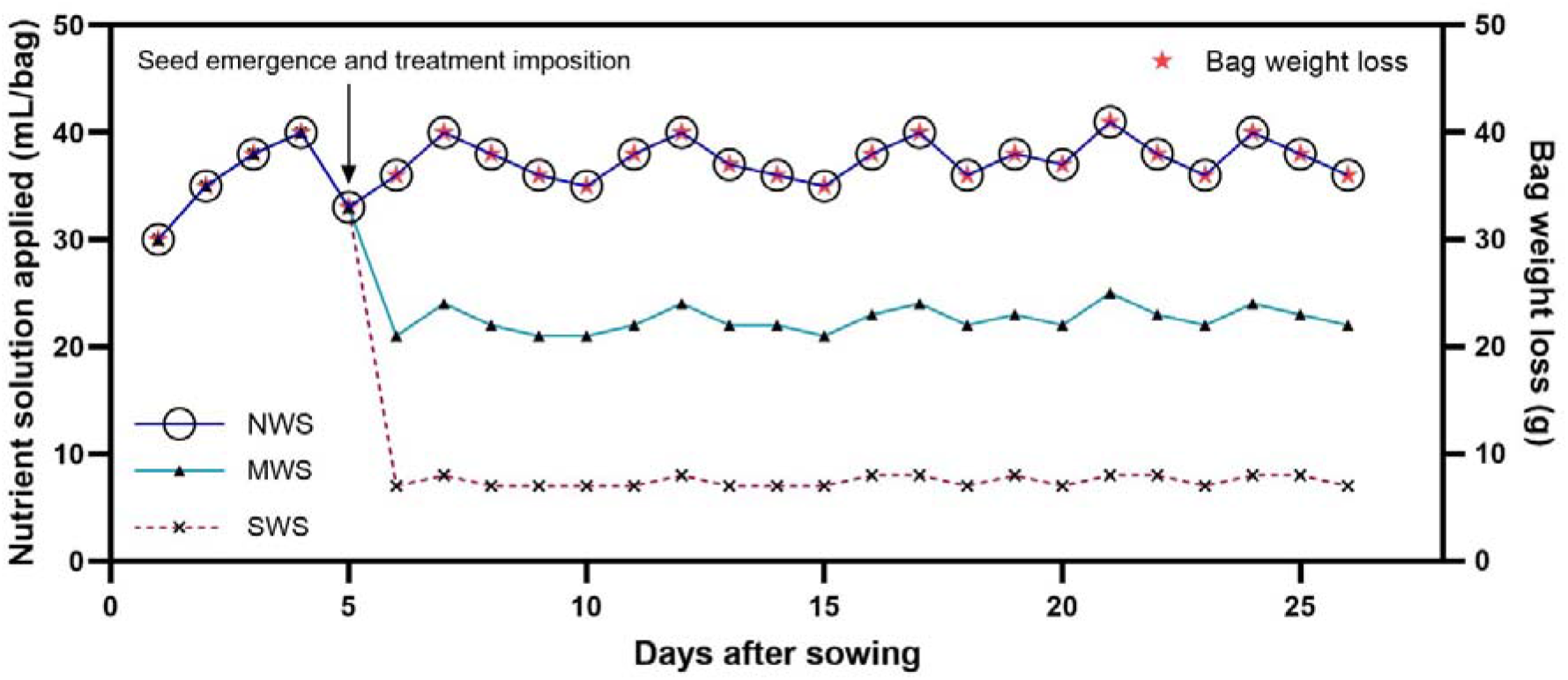
Daily weight loss of bag (g) and amount of nutrient solution applied (mL/bag) in different water stress treatments.

The Yoshida solution was modified to create three different K levels, including K- (Yoshida solution of 50% low K), K0 (Yoshida solution of normal K), and K+ (Yoshida solution of 50% high K), corresponding to 20 ppm K (35.7 g/L K_2_SO_4_), 40 ppm K (71.4 g/L K_2_SO_4_), and 60 ppm K (107.1 g/L K_2_SO_4_), respectively. The sulfur concentration was maintained by adjusting the amount of H_2_SO_4_ (Sp. gravity: 1.84, purity: 98%) according to K_2_SO_4_ concentrations in treatments (50 mL/L H_2_SO_4_ in K0, 61.14 mL/L in K−, and 38.85 mL/L in K+). All other nutrient concentrations of the solution remained unchanged. The pH of the solution was adjusted to 5.5 before the application.

A total of 405 root samples representing root architecture from five seedlings per treatment at three different time intervals (7, 14, and 21 days after emergence (DAE)) with three replications were analyzed. Roots were immersed underwater in a scanner acrylic tray (23.1×15.5×7 cm^3^) for 45 min before the scanning and stained with natural red dye (0.25 g/L) to increase contrast and reduce scanner light diffraction from the root surface. Root images were captured using an HP Scanjet G2410 scanner at a resolution of 600 dpi. A circle scale marker of 40 mm diameter was placed beneath the scanner tray, and all images were captured with the scale marker (Figure 1 C). Root samples collected at 21 DAE were used to estimate root dry weight; they were kept in the hot air oven at 65 °C ±1 °C overnight till constant weight was achieved.

Root architectural traits (Figure 3) were computed by digital imaging of root traits (DIRT)^14^. All scanned images were calibrated at the masking threshold level of 3 before computation (Figure 1 D and E). The output datasets (in pixels) were converted in metric units, except average root density (calculated as the ratio of foreground to background pixels within the root shape), using the following formula:

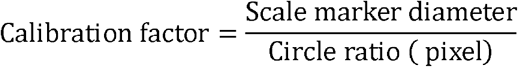

Where the circle ratio (in pixel) is computed by the DIRT platform.

*Root depth (mm)* = *l*ength in pixel x calibration factor

*Maximum width (mm)* = maximum width in pixel x calibration factor

*Projected root area (mm^2^)* = area in pixel x *calibartion factor*^2^

**Figure 3.**
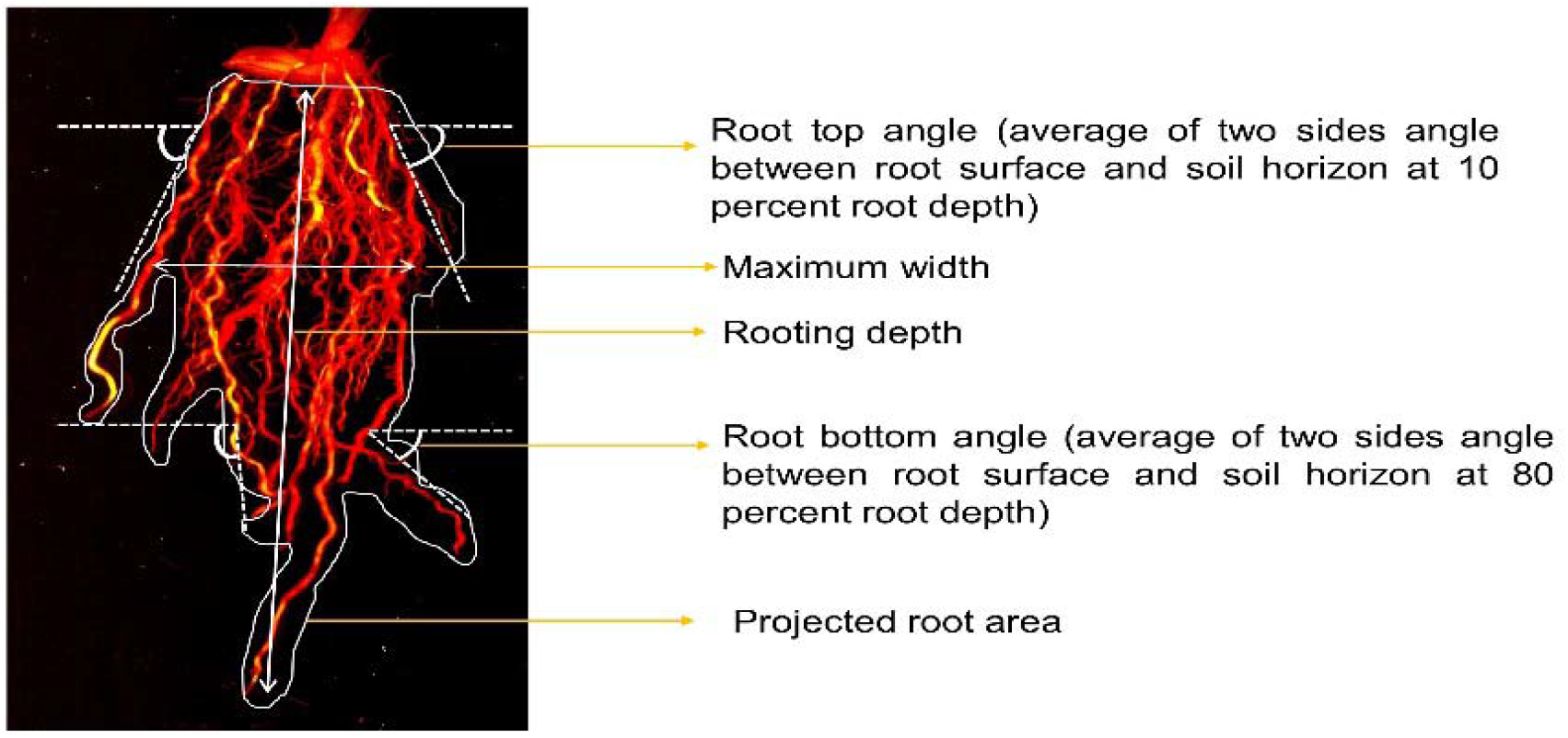
**R**ice seedling root traits accessed during the study

Shoot length (cm) and shoot dry weight (g) of the same seedlings from each treatment used for root analysis were measured at 21 DAE. Mean values of a dataset were subjected to analysis of variance (ANOVA)^15^ and the differences between means were considered to be statistically significant at p ≤ 0.05.

## RESULTS

### Vertical root architectural features

The projected root area (PRA) (Figure 4 A) decreased with increasing water stress, and the reduction was larger when the K level was also low. The PRA was reduced by 51% and 66% in PRA under mild (MWS) and severe water stresses (SWS) with low K level (K−), respectively, compared to non-stressed seedlings (NWS) treated with normal K level (K0). Additionally, a 25% reduction in PRA was observed under the same water stresses treatments (K− MWS and K− SWS) compared to those with normal K supply. However, higher K availability (K+) under MWS and SWS did not significantly influence the root area compared to the same water stress treatments, with normal K level.

**Figure 4.**
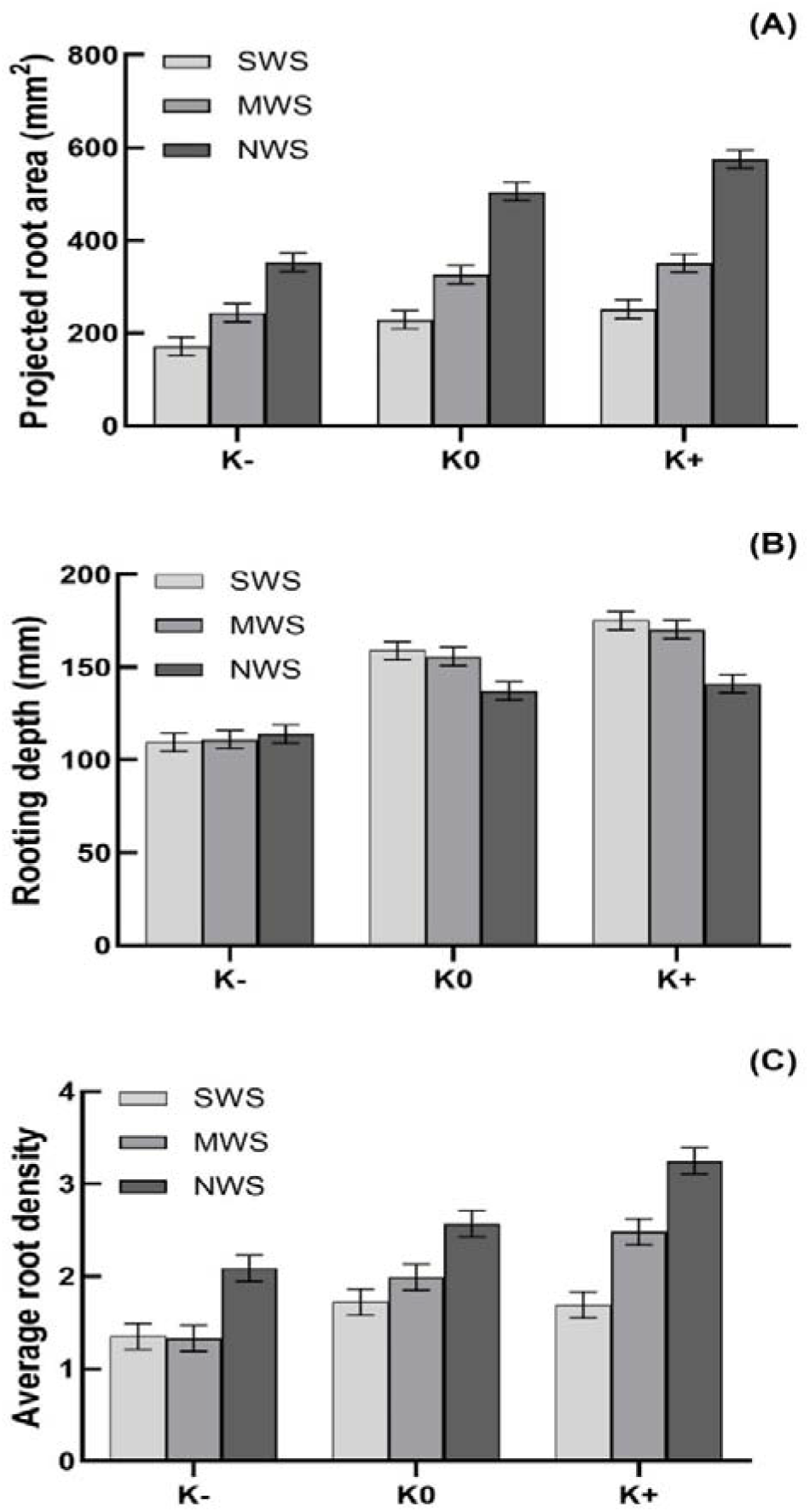
Projected root area (A), rooting depth (B) and average root density (C) at 21 DAE. NWS (no water stress), MWS (mild water stress) and SWS (severe water stress), K- (low potassium), K0 (normal potassium), K+ (high potassium) (mean ± S. Em.).

Seedling rooting depth (RD) (Figure 4 B) increased by water stress, though root depth decreased up to 28.7% and 31.1% under mild and severe water stresses with low K supply, respectively, as compared to the same water stress treatments with normal K availability. However, in contrast to low K, higher K availability increased root depth by 9.3% and 10% in MWS and SWS treatments over normal K supply, respectively. However, the effect of K+ on root depth on non-stressed treatments was not strong.

Average root density (Figure 4 C) was reduced with increasing water stress. Under mild water stress, K limitation reduced root density by 33% compared to the same water stress treatment with normal K supply. However, the application of a higher K level under MWS resulted in similar root density values as in non-stressed treatment with normal K level. There were no significant differences in root density between severe water stress treatments with low or high K supply, and was statistically similar to that under the same water stress with normal K supply.

### Horizontal root architectural features

The maximum width (MW) (Figure 5 A) of the root system decreased with increasing water stress. There was a larger decrease in root width with lower K supply, and 41% and 43% reductions in root width were recorded under MWS and SWS with low K level compared to the same water stress treatments with normal K supply, respectively. However, seedlings treated with higher K level under mild and severe water stress showed 20% and 48% higher root width values, compared to their counterparts with normal K supply, respectively.

**Figure 5.**
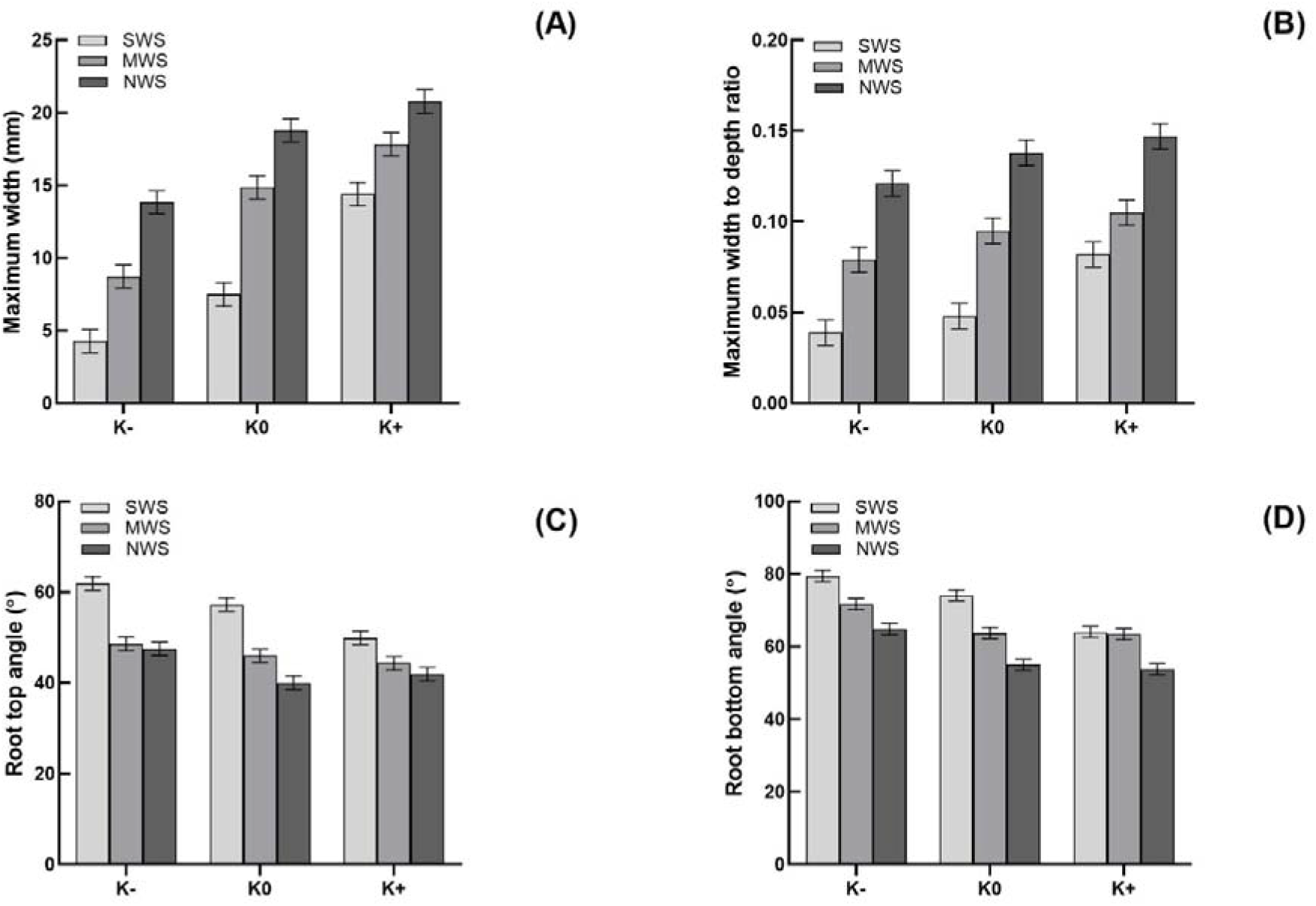
Maximum width (A), maximum width to rooting depth ratio (B), root top angle (C) and root bottom angle (D) at 21 DAE. NWS (no water stress), MWS (mild water stress), SWS (severe water stress), K- (low potassium), K0 (normal potassium), K+ (high potassium) (mean ± S. Em.).

Water stress reduced the MW/RD ratio (Figure 5 B). The increased K availability (K+) increased the MW/RD ratio by 6.5%, while low K availability (K−) decreased it by 12.3% over the normal availability (K0) in non-stressed seedlings. Although under different water regimes, the availability of K did not influence the MW/RD ratio at 21 DAE, their interaction was significant at 7 and 14 DAE (Supplementary Figure S2 A & B).

Narrower root top angle (RTA) (Figure 5 C) was found in non-stressed treatment, while the widest RTA was obtained in roots under severe water stress. The RTA was increased by 6° and 17.2° under mild and severe water stress with normal K supply compared to non-stressed treatment, respectively. Under MWS, different levels of K had almost similar effects on RTA. However, rice seedlings exposed to severe water stress with higher K supply showed narrower RTA (49.9°) as compared to their counterpart with low K supply (61.9°), with the value almost similar to that in mild water stress treatment with normal K supply (46.0°).

Similar to RTA, water stress increased the root bottom angle (RBA) (Figure 5 D). The RBA values were 63.7° and 74.1° in mild and severe water stress treatments with normal K level, which increased by 8.7° and 19.1° compared to non-stressed condition, respectively. The RBA was wider under water stress with a low K level. Rice seedlings subjected to MWS and SWS with low K availability showed RBA values of 71.7° and 79.4°, which were 8° and 5.3° higher than their counterparts with normal K supply, respectively. The RBA in seedlings treated with K+ under MWS did not show any difference with that with normal K supply, though under SWS condition, higher K supply reduced RBA by 10°.

### Root dry weight

Root dry weight (RDW) (Figure 6 B) decreased with increasing water stress. The RDW of rice seedlings grown under low K condition and exposed to MWS and SWS showed 31% and 49% reductions as compared to their counterparts with normal K availability (K0), respectively. However, the application of high K to rice seedlings grown under severe water stress resulted in a 15.6% increase in RDW over their counterpart with normal K supply (K0); however, such an increase was not observed in seedlings subjected to MWS.

**Figure 6.**
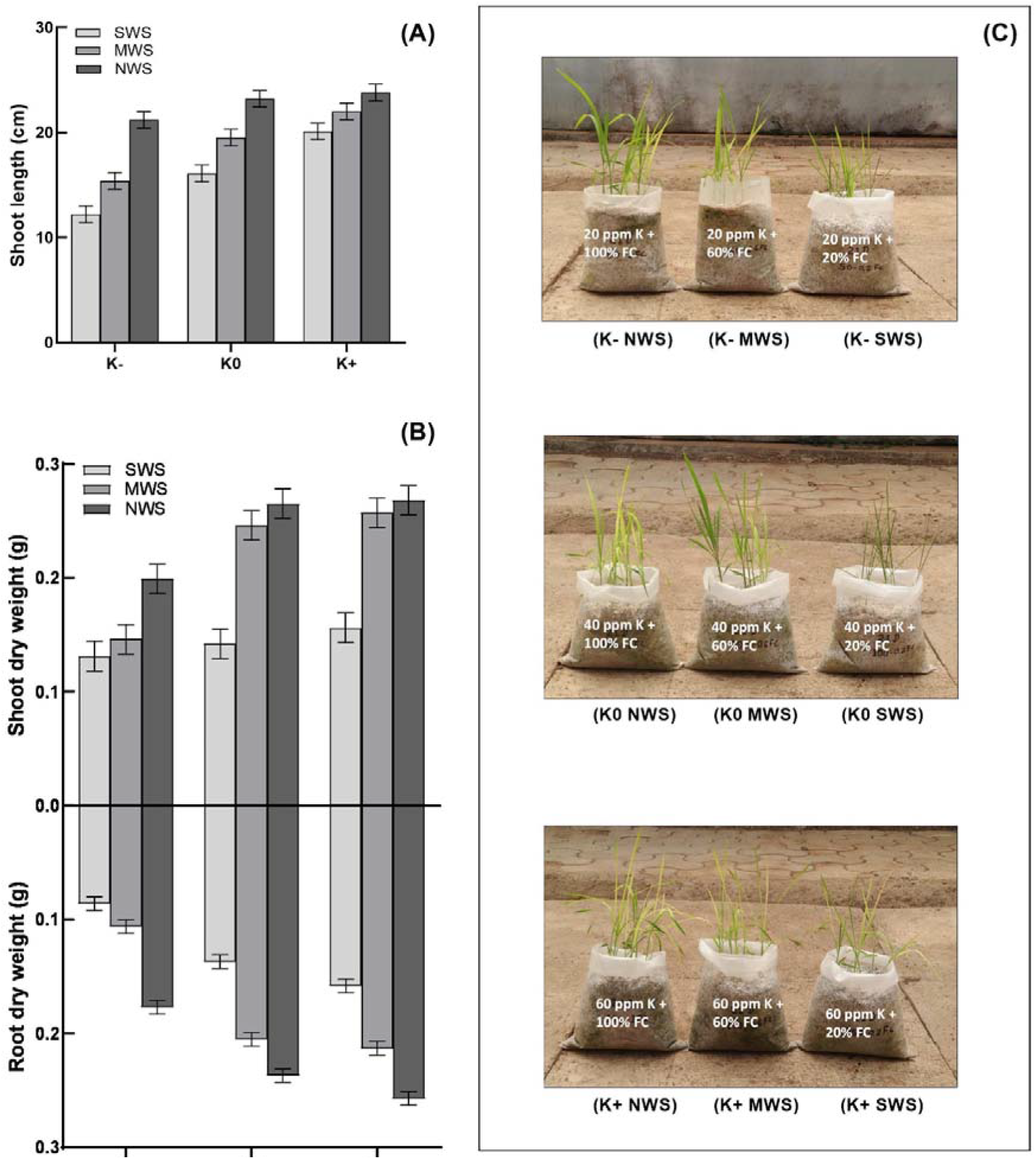
Shoot length (A), shoot and root dry weight (B) and rice seedlings growth (C) at 21 DAE. NWS (no water stress), MWS (mild water stress), SWS (severe water stress), K- (low potassium), K0 (normal potassium), K+ (high potassium) (mean ± S. Em.).

### Seedling growth

As shown in Figure 6 A, the shoot lengths of rice seedlings treated with K0 under mild and severe water stress conditions were 16% and 30% lower than that of non-stressed seedlings, respectively. Shoot length in seedlings treated with low and high K levels under non-stressed conditions did not show any significant differences. However, seedlings exposed to MWS and SWS with low K level showed 21% and 24% lower shoot lengths as compared to those grown with normal K supply under the same water stress conditions, respectively. High K application to rice seedlings grown under mild and severe water stress conditions increased shoot length by 12.8% and 24.8% as compared to that of their counterparts treated with normal K availability, respectively. Shoot dry weight of seedlings (Figure 6 B) decreased with increasing water stress. Such reduction was larger by 40% in rice seedlings receiving low K level under MWS as compared to those grown under the same moisture stresses with normal K supply while seedlings subjected to mild water stress with high K level had similar shoot dry weight values as those receiving normal K supply. However, under SWS, low and high K does not show any significant effect. Similarly, total dry weight (Figure S3) of seedlings subjected to mild and severe water stress with high K level had similar total dry weight values as those receiving normal K supply while seedlings grown with low K under mild and severe water stress had shown significant reduction in total dry weight.

## DISCUSSION

Although root architecture is regulated by the genetic makeup of the variety, it shows a wide range of phenotypic plasticity in response to nutrient^16^ and water availability^17^, and plays a crucial role in the tolerance of rice plant to drought stress^12^. In the present study, the root architecture of rice seedlings was found to be shaped differently by water and K availability (Figure 7 and Figure 8). The projected root area and average root density decreased with increasing water stress, and this negative effect was stronger with lower K levels, while such reduction was not observed with normal and high levels of K under mild and severe water stresses. Similarly, in tomato, K increases root surface area, volume and number of root tips under atmospheric drought^18^. The more profound effect of water stress on these parameters indicates that both K and water were limiting factors, which may regulate root features independently. Sugar translocation is dependent on K availability^19^ and triggers cell division^20^, while cell elongation is dependent on the level of cell turgor^21^ and water content. Our study indicates that the sufficient availability of K is more important for root system development, as non-stressed rice roots receiving low K had reduced root area compared to their counterpart seedlings grown with normal K supply, which may be due to the lack of turgor pressure as a result of low water absorption^21^ and/or disruption of auxin maxima in the root tip under low K conditions^22^.

**Figure 7.**
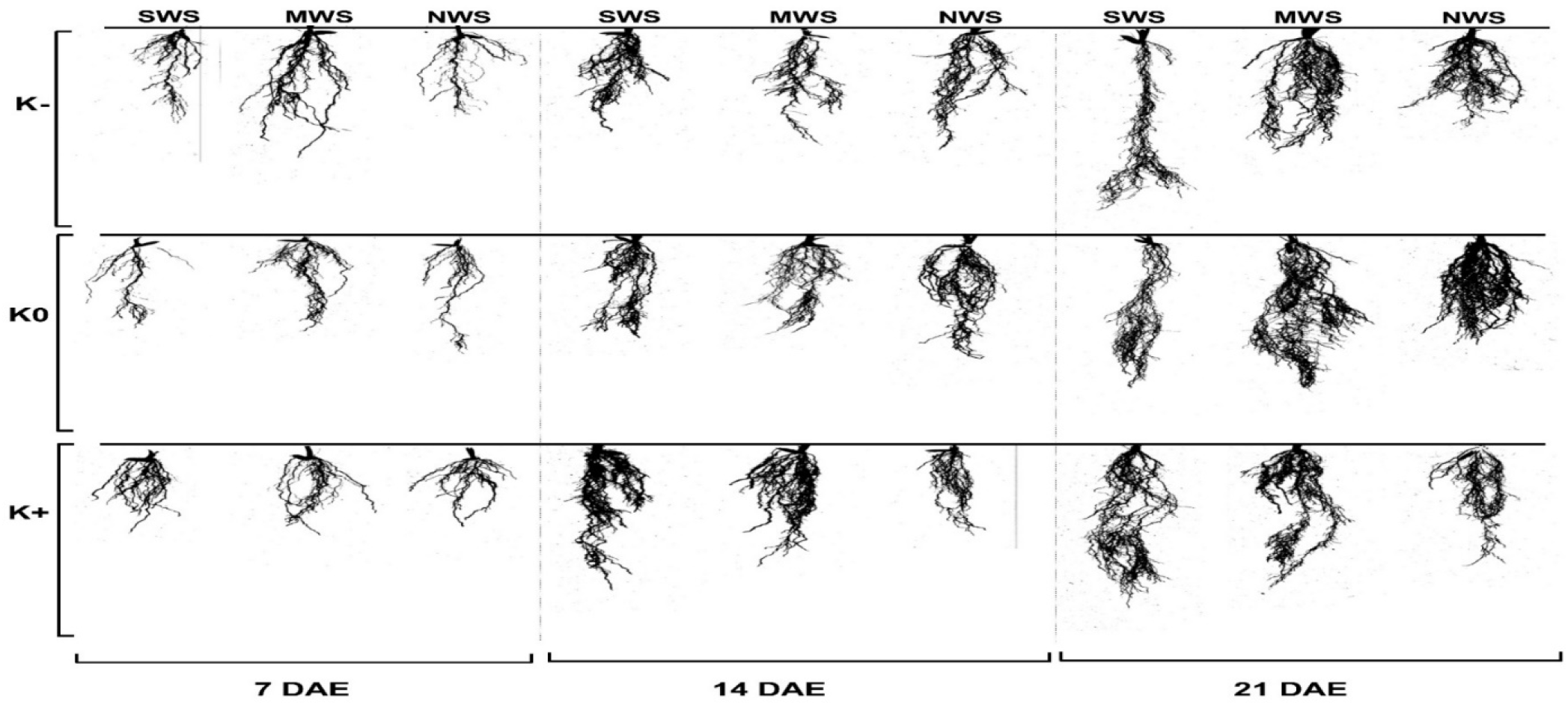
Rice root architecture under SWS (severe water stress), MWS (mild water stress) and NWS (no water stress) with K- (low potassium), K0 (normal potassium) and K+ (high potassium) at 7, 14 and 21 DAE.

**Figure 8.**
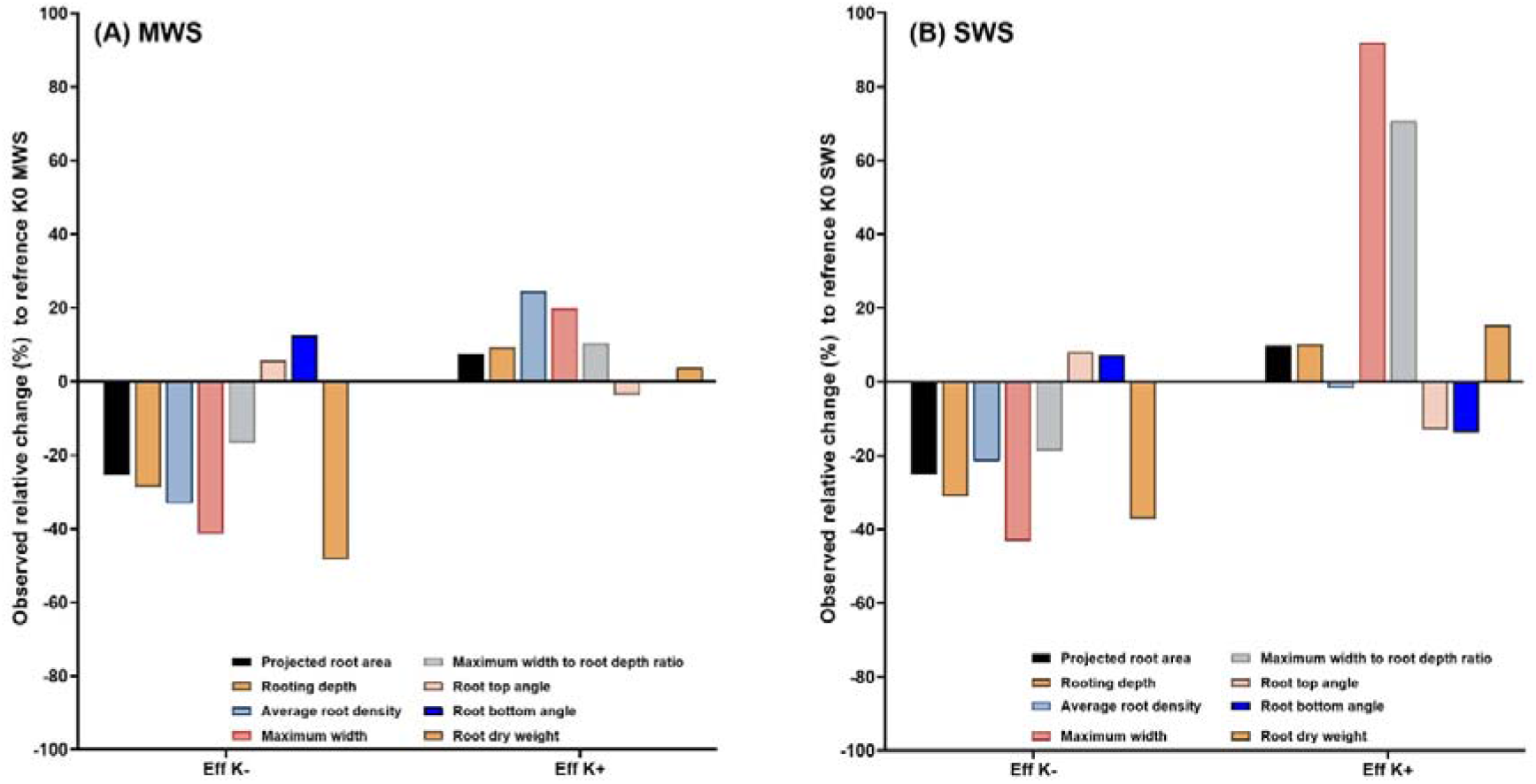
Observed relative changes (%) in root traits due to effect of K- (low potassium) and K+ (high potassium) under MWS (mild water stress) (A) and SWS (severe water stress) (B). Individual effect was calculated over K0 (normal potassium) with respective water stress.

The acquisition of soil resources is dependent on the horizontal and vertical expansion of the root. In the present study, to detect a preferential direction of horizontal and vertical expansion of roots, root depth, maximum width, and the ratio of the maximum width to root depth were estimated. The root depth was increased and the maximum width was decreased with increasing water stress, which resulted in the lower MW/RD ratio, indicating that under stress, vertical elongation of root is stimulated over horizontal, to seek the water source deeper down in the soil, thus steeper root growth occurs. The addition of K under water stress promoted horizontal elongation by increasing root width and increasing MW/RD. Moreover, roots with steep growth also have wide root top and bottom angles under MWS and SWS, indicating that roots had lower surface areas of contact. In field condition, this will limit water and nutrient uptake from the soil. High availability of K in seedlings under MWS resulted in compensation for the reduction in the top and bottom angles, but under SWS, high K application failed to exert this positive effect. This study suggests that besides increasing translocation of mineral nutrients in xylem by K^23^ under water stress, K fertilization helps in maintaining RTA and RBA, and thus increases root-soil contact area and may contribute in nutrient uptake by exploring more soil areas. Auxin synthesis, distribution, and signaling are the principal aspects of rice root development^24^. The depth of the root system is controlled by an auxin-inducible DRO gene, which regulates root top angle determining the direction of root elongation^2^. However, the mechanism of observed changes in root top and bottom angles by K availability is unclear, though K found to influence auxin signaling and distribution, as in rice, K transporter alters the membrane-bound auxin efflux proteins^25, 26^ and may induces root angle changes.

Seedling growth reduces due to water stress and reduction in root dry weight, shoot dry weight and seedling height was more with low potassium. However, high K alleviate these effects and increased shoot length, shoot dry weight and root dry weight, though the beneficial effect of high K was distinct in mild water stress. High K application under severe water stress does not increase gain in total dry weight over normal potassium. These results are in accordance with earlier study^27^.

## CONCLUSION

This study shows that the interaction of K and WS influenced root spatial distribution, which could offer the advantage of exploring underground resources for promoting above-ground growth of rice seedlings under mild water stress, depending upon the K availability. Our results emphasize the necessity to maintain the potassium status of soils under moisture deficit conditions. These findings extend our knowledge on the role of K in drought tolerance through root architecture modification. Future studies should aim to evaluate the extent to which the root architecture responds to water stress and K availability in broad genetic background.

**Table 1.** Summary of analysis of variance for water stress (WS), potassium (K) and their interactions (WS x K) on root architectural characteristics and seedling growth.

| Parameters<sup>†</sup> | p Values |  |  |
| --- | --- | --- | --- |
|  | WS | K | WS x K |
| Projected root area (mm<sup>2</sup>) at 7 DAE | * | * | ns |
| Projected root area (mm<sup>2</sup>) at 14 DAE | * | * | ns |
| Projected root area (mm<sup>2</sup>) at 21 DAE | * | * | * |
| Rooting depth (mm) at 7 DAE | * | * | ns |
| Rooting depth (mm) at 14 DAE | * | * | * |
| Rooting depth (mm) at 21 DAE | * | * | * |
| Maximum width (mm) at 7 DAE | * | * | ns |
| Maximum width (mm) at 14 DAE | * | * | ns |
| Maximum width (mm) at 21 DAE | * | * | * |
| Average root density at 7 DAE | * | * | ns |
| Average root density at 14 DAE | * | * | * |
| Average root density at 21 DAE | * | * | * |
| Maximum width to depth ratio at 7 DAE | * | * | * |
| Maximum width to depth ratio at 14 DAE | * | * | * |
| Maximum width to depth ratio at 21 DAE | * | * | ns |
| Root top angle (°) at 7 DAE | * | * | ns |
| Root top angle (°) at 14 DAE | * | * | ns |
| Root top angle (°) at 21 DAE | * | * | * |
| Root bottom angle (°) at 7 DAE | * | * | ns |
| Root bottom angle (°) at 14 DAE | * | * | ns |
| Root bottom angle (°) at 21 DAE | * | * | * |
| Root dry weight (g) | * | * | * |
| Shoot length (cm) | * | * | * |
| Shoot dry weight (g) | * | * | * |
| Total dry weight (g) | * | * | * |

<sup>†</sup>Root traits of 7 and 14 DAE (days after emergence) are presented in supplementary figures: Figure S1 to S2.
\*Significant at $p \geq 0.05$ , ns: not significant.

## REFERENCES

1. Gowda V. R. P., Henry A., Yamauchi A., Shashidhar H. E. and Serraj R., Root biology and genetic improvement for drought avoidance in rice. Field Crop Res., 2011, 122, 1–13.

2. Uga Y., Sugimoto K., Ogawa S., Rane J., Ishitani M., Hara N., Control of root system architecture by DEEPER ROOTING 1 increases rice yield under drought conditions. Nat. Gen., 2013, 45, 1097–1102.

3. Kudoyarova G.R., Dodd I.C., Veselov D.S., Rothwell S.A., Yu. Veselov S., Common and specific responses to availability of mineral nutrients and water. J. Exp. Bot., 2015, 66, 2133–2144.

4. Grzebisz W., Gransee A., Szczepaniak W. and Diatta J., The effects of potassium fertilization on water use efficiency in crop plants. J. Plant Nutr. Soil Sci., 2013, 176, 355–374.

5. Zain N.A. M. and Ismail M.R., Effects of potassium rates and types on growth, leaf gas exchange and biochemical changes in rice (Oryza sativa) planted under cyclic water stress. Agricultural Water Management, 2016, 164, 83–90.

6. Filho A.C. A.C., Crusciol C.A. C., Nascente A.S., Mauad M. and Garcia R.A., Influence of potassium levels on root growth and nutrient uptake of upland rice cultivars. Rev. Caatinga, 2017, 30 (1), 32–44

7. Tatsumi J., Endo N. and Kono Y., Root growth and partitioning of 13C-labelled photosynthate in the seminal root of corn seedlings as affected by light intensity. Jpn. J. Crop Sci., 1992, 61, 271–278.

8. Mengel K. and Arneke W.W., Effect of potassium on the water potential, the pressure potential, the osmotic potential and cell elongation in leaves of Phaseolus vulgaris. Physiol. Plant, 1982, 54, 402–408.

9. Benlloch-González M., Arquero O.J., Fournier M., Barranco D. and Benlloch M., K+ starvation inhibits water-stress-induced stomatal closure. J. Plant Physiol., 2008, 165, 623–630.

10. Cakmak I., The role of potassium in alleviating detrimental effects of abiotic stresses in plants. J. Plant Nutr. Soil Sci., 2005, 168, 521–530.

11. Jordan-Meille L., Martineau E., Bornot Y., Lavres J., Abreu-Jr. H.C., and Domec J.C., How does water-stressed corn respond to potassium nutrition? A shoot-root scale approach study under controlled conditions. Agriculture, 2018, 8, 180.

12. Kano-Nakata M., Gowda V.R. P., Henry A., Serraj R., Inukai Y., Fujita D., Kobayashi N., Roel R. Suralta R.R. and Yamauchi A., Functional roles of the plasticity of root system development in biomass production and water uptake under rainfed lowland conditions. Field Crops Research, 2013, 144, 288–296.

13. Yoshida S., Forno D.A., Cock J.H. and Gomez K.A., Laboratory Manual for Physiological Studies of Rice. (2nd edition). The International Rice Research Institute, Philippines, 1976.

14. Das A., Schneider H., Burridge J., Ascanio A.K. M., Wojciechowski T., Topp. C.N., Lynch P.J., Weitz J.S. and Bucksch A., Digital imaging of root traits (DIRT): a high throughput computing and collaboration platform for field-based root phenomics. Plant Methods. 2015, 11, 51.

15. Gomez K.A. and Gomez A.A., Statistical procedures for agricultural research (2 ed.). John wiley and sons, New York, 1984.

16. Shahzad Z. and Amtmann A., Food for thought: how nutrients regulate root system architecture. Current Opinion in Plant Biology, 2017, 39, 80–87.

17. Giehl R.F. H. and Von Wiren N., Hydropatterning-how roots test the waters. Science, 2018, 362, 1358–1359.

18. Zhang J., Jiao X., Du Q., Song X., Ding J. and Li J., Effects of vapor pressure deficit and potassium supply on root morphology, potassium uptake, and biomass allocation of tomato seedlings. J Plant Growth Regul., 2020, https://doi.org/10.1007/s00344-020-10115-2

19. Martineau E., Domec J.C., Bosc A., Dannoura M., Gibon Y., Bernard C., Jordan-Meille L., The role of potassium on maize leaf carbon exportation under drought condition. Acta Physiol. Plant, 2017, 39, 219, 1–13.

20. Wang L., & Ruan Y.L., Regulation of cell division and expansion by sugar and auxin signaling. Frontiers in plant science, 2013, 4, 163, 1–9.

21. Gerardeaux E., Jordan-Meille L., Constantin J., Pellerin S. and Dingkuhn M., Changes in plant morphology and dry matter partitioning caused by potassium deficiency in Gossypium hirsutum (L.). Environ. Exp. Bot., 2010, 67, 451–459.

22. Song W., Liu S., Meng L., Xue R., Wang C., Liu G., Dong C., Wang S., Dong J. and Zhang Y., Potassium deficiency inhibits lateral root development in tobacco seedlings by changing auxin distribution. Plant Soil, 2015, 396 (1/2), 163–173.

23. Hasanuzzaman M., Borhannuddin B.M. H. M., Nahar K., Hossain Md. S., Mahmud Al J., and Hossen Md. S., Masud C.A. A., Moumita and Fujita M., Potassium: A vital regulator of plant responses and tolerance to abiotic stresses. Agronomy, 2018, 8, 31, 1–29.

24. Wang Y., Zhang T., Wang R., and Zhao Y., Recent advances in auxin research in rice and their implications for crop improvement, Journal of Experimental Botany, 2018, 69, 255–263.

25. Yang T., Feng H., Zhang S., Xiao H., Hu Q., Chen G., Xuan X., Moran N., Murphy A., Yu L. and Xu G. The potassium transporter OsHAK5 alters rice architecture via ATP-dependent transmembrane auxin fluxes. Plant Commun., 2020, 1, 100052, 1–15.

26. Sustr M., Soukup A. and Tylova E. Potassium in root growth and development. Plants, 2019, 8, 435, 1–16.

27. Zain N.A. and Ismail M.R. Effects of potassium rates and types on growth, leaf gas exchange and biochemical changes in rice (Oryza sativa) planted under cyclic water stress. Agricultural Water Management. 2016, 164, 83–90.

