## Supplementary figures and images for "Does plant root architecture respond to potassium nutrition under water stress? A case from rice seedling root responses"

### S1

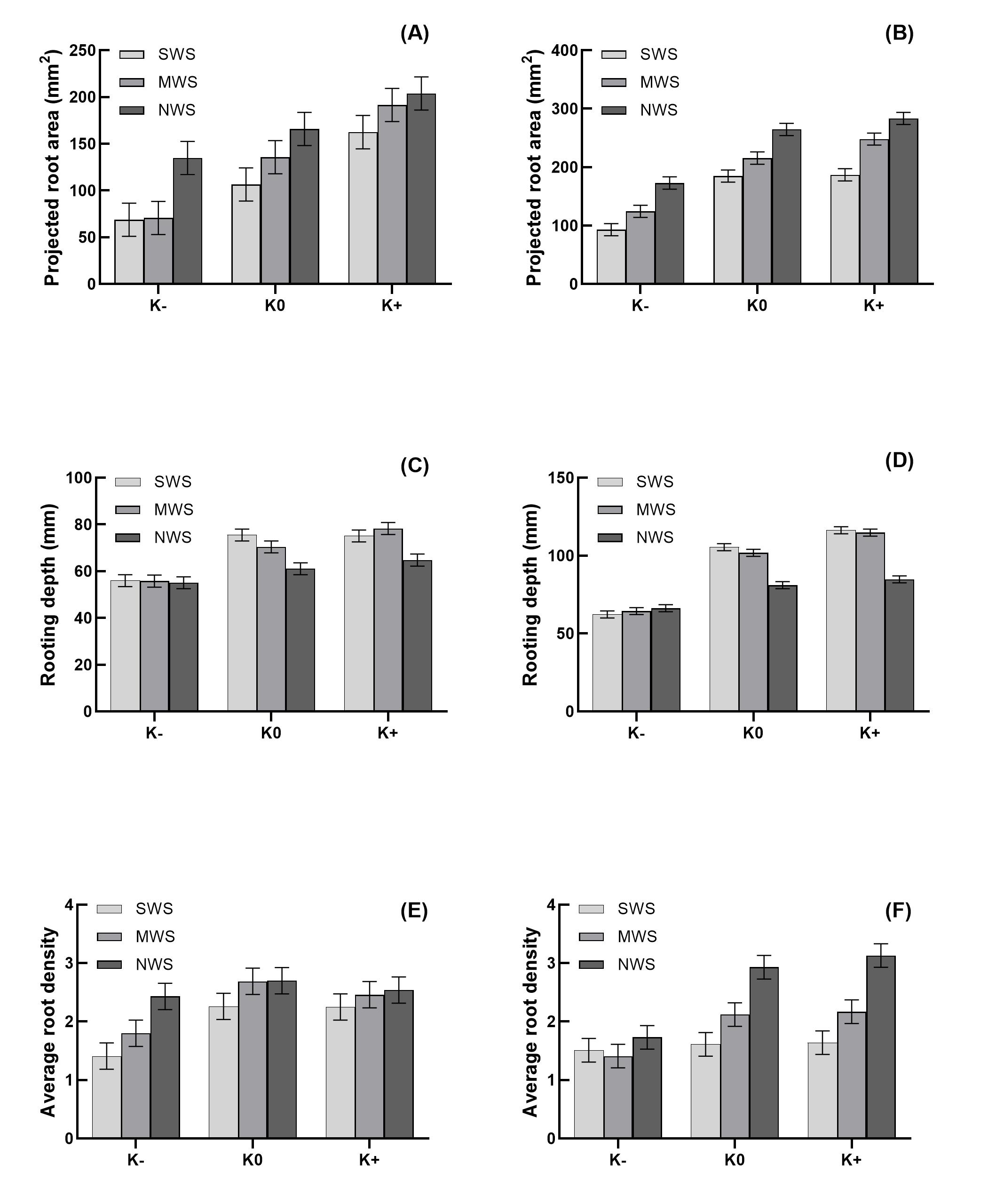

### S2

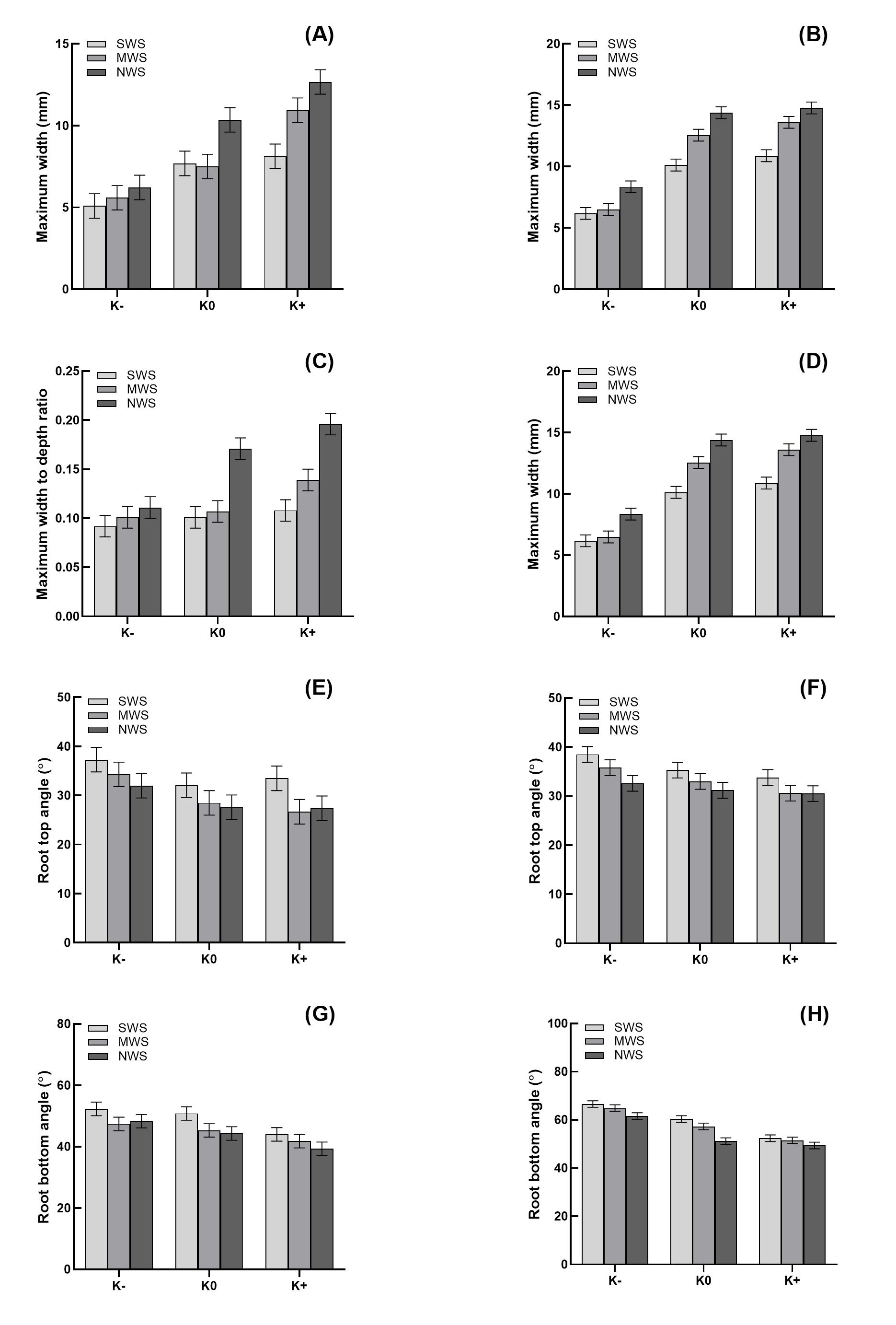

### S3

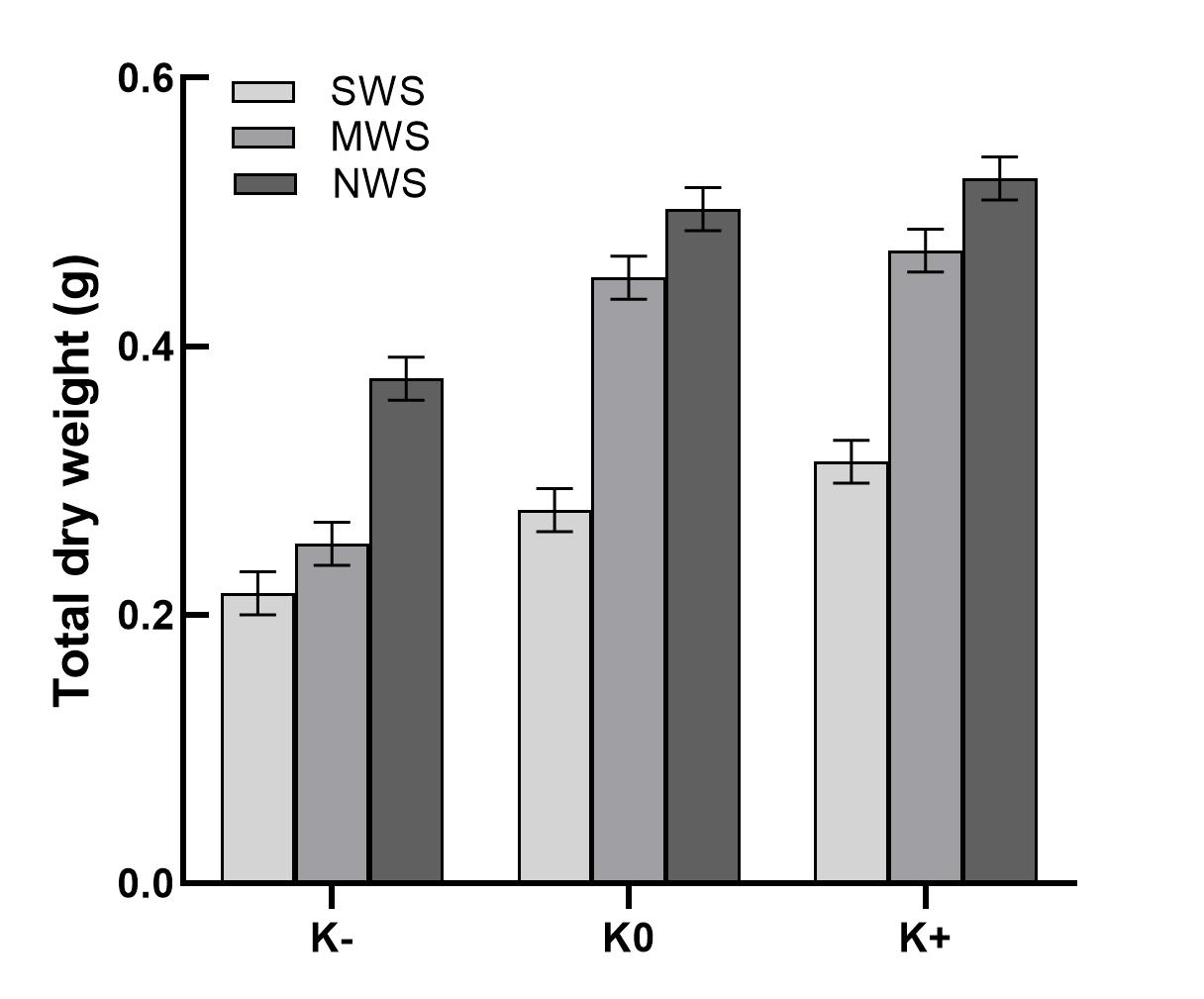
